# Structural imaging of native cryo-preserved secondary cell walls reveals presence of macrofibrils composed of cellulose, lignin and xylan

**DOI:** 10.1101/648360

**Authors:** Jan J. Lyczakowski, Matthieu Bourdon, Oliver M. Terrett, Ykä Helariutta, Raymond Wightman, Paul Dupree

## Abstract

The woody secondary cell walls of plants are the largest repository of renewable carbon biopolymers on the planet. These walls are made principally from cellulose and hemicelluloses and are impregnated with lignin. Despite their importance as the main load bearing structure for plant growth, as well as their industrial importance as both a material and energy source, the precise arrangement of these constituents within the cell wall is not yet fully understood. We have adapted low temperature scanning electron microscopy (cryo-SEM) for imaging the nanoscale architecture of angiosperm and gymnosperm cell walls in their native hydrated state. Our work confirms that cell wall macrofibrils, cylindrical structures with a diameter exceeding 10 nm, are a common feature of the native hardwood and softwood samples. We have observed these same structures in *Arabidopsis thaliana* secondary cell walls, enabling macrofibrils to be compared between mutant lines that are perturbed in cellulose, hemicellulose and lignin formation. Our analysis indicates that the macrofibrils in Arabidopsis cell walls are composed, at least partially, of cellulose, xylan and lignin. This study is a useful additional approach for investigating the native nanoscale architecture and composition of hardwood and softwood secondary cell walls and demonstrates the applicability of Arabidopsis genetic resources to relate fibril structure with wall composition and biosynthesis.

## Introduction

The majority of carbon in terrestrial biomass is stored in forests as wood (Ramage et al., 2017, Pan et al., 2011). The current classification system distinguishes two types of timber. Wood from Angiosperm trees is known as hardwood and the wood made by Gymnosperm species is described as softwood (Ramage et al., 2017). Despite significant differences in tissue organisation and chemical composition, both these types of timber are almost entirely formed from plant secondary cell walls – an extracellular matrix made primarily from cellulose, lignin and hemicelluloses (Schweingruber, 2007). Considering the ecological and industrial importance of wood and other cell wall materials, our knowledge of the exact arrangement of these polymers in the cell wall remains poor. A better understanding of the molecular architecture and ultrastructure of cell walls is needed to describe the complex spatiotemporal deposition pattern of the cell wall polymers. This may contribute to the development of more efficient biofuel feedstocks (Loque et al., 2015), to the improvement in our understanding of novel biomaterials such as nanocellulose (Jarvis, 2018), and to applications such as advanced approaches for the use of timber in the construction industry (Ramage et al., 2017)

Cellulose is the main constituent of plant cell walls (Pauly and Keegstra, 2008). At the molecular level, cellulose has a simple repeating structure of β-1,4-linked glucopyranosyl residues. These glucan chains coalesce to form a crystalline cellulose microfibril. The exact structure of the microfibril is unknown, however, it has been suggested the elementary microfibril consists of 18 or 24 individual glucan chains (Gonneau et al., 2014, Hill et al., 2014, Turner and Kumar, 2017). Individual cellulose microfibrils associate to form larger order structures known as macrofibrils (Niklas, 2004). In plant primary cell walls this close-contact association may be limited to selected parts of microfibril which is proposed to lead to formation of so-called biomechanical hotspots (Cosgrove, 2014). A range of imaging and spectroscopic techniques has been used to investigate cellulose macrofibrils in secondary cell walls, as reviewed by (Purbasha et al., 2009), but due to technical challenges the precise structure in native, unprocessed, hydrated secondary cell walls remains poorly described. Lignin is the main non-polysaccharide component of both hardwood and softwood and is made by coupling of monolignol radicals in secondary cell walls. Three main monolignols exist in plants, which, once turned into chemical radicals by the activity of laccases and peroxidases, can couple in a random manner to form a lignin polymer made from guaiacyl (G), syringyl (S), and p-hydroxyphenyl (H) units (Ralph et al., 2004). The monolignol composition of hardwood and softwood differs, with the former consisting of predominantly S and G units and the latter being made almost solely from G units (Vanholme et al., 2010). The process of lignification is important for wood mechanical properties. Arabidopsis mutant plants with reduced lignin content or altered monolignol composition often have collapsed xylem vessels and can be severely dwarfed (Bonawitz and Chapple, 2010). Lignin is proposed to associate with cell wall polysaccharides to form the recalcitrant matrix (Terrett and Dupree, 2019).

Xylan and galactoglucomannan are the principal hemicelluloses in hardwood and softwood. Xylan is a polymer of β-1,4-linked xylopyranosyl residues and is the main hemicellulose in hardwood but is also present in softwood (Scheller and Ulvskov, 2010). Hardwood and softwood xylans carry α-1–2 linked glucuronic acid (GlcA) branches which can be methylated on carbon 4 leading to formation of 4-O-Methylglucuronic acid (MeGlcA) (Scheller and Ulvskov, 2010). In addition to GlcA and Me GlcA (together, [Me]GlcA) decorations, hardwood xylan hydroxyls are acetylated on carbon 2, carbon 3 or both carbons of the monomer. The softwood xylan, in addition to the MeGlcA branches, carries α-1,3–linked arabinofuranosyl decorations (Scheller and Ulvskov, 2010, Busse-Wicher et al., 2016b). The presence of [Me]GlcA branches on xylan is important for the maintenance of biomass recalcitrance (Lyczakowski et al., 2017) and, together with acetylation in hardwood and arabinose decorations in softwood, these substitutions are mostly distributed with an even pattern on xylosyl units (Bromley et al., 2013, Busse-Wicher et al., 2014, Busse-Wicher et al., 2016b, Martinez-Abad et al., 2017). This so-called ‘compatible’ patterning of xylan substitutions is thought to allow the hydrogen bonding between xylan, in a two-fold screw conformation, and the hydrophilic surface of the cellulose microfibril (BusseWicher et al., 2016a, Simmons et al., 2016, Grantham et al., 2017). Galactoglucomannan (GGM) is the main hemicellulose in softwood (Scheller and Ulvskov, 2010) but is also present in hardwood xylem. GGM has a backbone formed from both β-1,4-linked mannosyl and glucosyl residues with some mannosyl residues substituted by an α-1,6-linked galactosyl branch. The GGM backbone can also be acetylated. The arrangement of mannose and glucose units in softwood GGM is thought to be random, but a recently described regular structure GGM found in Arabidopsis mucilage was proposed to bind to both the hydrophilic and hydrophobic surface of the cellulose microfibril (Yu et al., 2018). *In vitro* studies using TEM and 1D ^13^C NMR indicate that a range of branched and unbranched mannan and glucomannan structures can interact with bacterial cellulose (Whitney et al., 1998). Softwood GGM is also proposed to interact with the cellulose microfibril (Terashima et al., 2009) and recent evidence demonstrates that it can form covalent linkages with lignin (Nishimura et al., 2018).

Although we now have a better understanding of secondary cell wall composition and the nature of the interactions between its main constituents, a picture of the ultrastructural assembly of wall polymers into a secondary cell wall matrix is not yet complete. Solid state NMR (ssNMR) analysis has been applied extensively to the study of polymer interactions in both primary and secondary walls. This, for example, provided evidence that in dried primary wall samples from Arabidopsis, pectin and xyloglucan may be interacting with the cellulose microfibril (Dick-Perez et al., 2011). Analysis of hydrated secondary cell wall of Arabidopsis with solid state NMR indicated that xylan is likely to interact with the hydrophilic surface of the cellulose microfibril as a two-fold screw (Simmons et al., 2016, Grantham et al., 2017). Recent ssNMR analysis indicates that in dried cell walls of grasses xylan is likely to interact with lignin (Kang et al., 2019). Despite providing excellent insights into the proximity of different cell wall components ssNMR cannot provide information about the assembly of these constituents into higher order structures. Some insights into this process have been achieved with other techniques. This includes application of vibrational microspectroscopy techniques such as FT-IR and RAMAN to study the orientation of cellulose and other cell wall components in the matrix, as reviewed by (Gierlinger, 2018). AFM has been applied to the study of cell wall matrix assembly, but the work has been focused on primary cell walls (Cosgrove, 2014) and only recent advances allowed nanoscale resolution imaging of dried spruce secondary cell walls (Casdorff et al., 2017). Moreover, insights into the assembly of cellulose microfibrils in wood walls of conifers (Fernandes et al., 2011) and dicots (Thomas et al., 2014) have been obtained using wide-angle X-ray scattering (WAXS) and small-angle neutron scattering (SANS).

In addition to these various approaches, other studies have attempted to use scanning electron microscopy (SEM) to study the structure of plant cell walls. Low temperature SEM (cryo-SEM), in which the sample is rapidly frozen and then maintained cold during imaging, has been used to study the collapse of pine needle tracheid cell walls upon prior dehydration (Cochard et al., 2004) and to visualise the bulging of root hairs in the *kojak* (cellulose synthase-like) mutant (Favery et al., 2001). Additionally, higher magnification cryo-SEM has been used to visualise cell walls of wheat awns (Elbaum et al., 2008). Some awn cell walls exhibit structural differences that are dependent upon the level of hydration and cryo-SEM revealed extensive layering within the wall, however, the technique was not further optimised to investigate individual fibrils. Field emission (FE) SEM techniques were effectively used to study the alignment of cellulose microfibrils in Arabidopsis hypocotyls (Refregier et al., 2004), roots (Himmelspach et al., 2003) and stems (Fujita et al., 2013). FE-SEM has also been applied to investigate wood structure, including observations of microfibril alignment in fixed cell walls of fir tracheids (Abe et al., 1997) and lignin distribution in spruce tracheids (Fromm et al., 2003). Importantly, FE-SEM analysis of dehydrated pine and poplar wood suggests that secondary cell walls of these species contain macrofibrils – cylindrical fibrillar structures with a diameter of up to 60 nm, which presumably comprise of bundles of elementary cellulose microfibrils (Donaldson, 2007). Moreover, the diameter of these macrofibrils was observed to increase with increasing lignification, suggesting that the macrofibrils may be formed from association of lignin and cell wall polysaccharides. This analysis was extended further to wood from Ginkgo where the FE-SEM was combined with density analysis to propose a model of macrofibril formation based on cellulose, GGM, xylan and lignin interaction (Terashima et al., 2009).

It has been suggested that some of the treatments used in preparation of the FE-SEM cell wall samples have little impact on the microfibril alignment and that the technique may provide a true representation of native (unprocessed) cell wall features (Marga et al., 2005). The FE-SEM techniques applied to secondary cell wall samples, however, included additional steps such as (i) fixation and exposure to organic solvents (ii) a thermal treatment that may result in some degree of wall degradation (Fromm et al., 2003) and (iii) a thick coating of heavy metal which may impact upon the resolution (Donaldson, 2007), raising questions about the effect these may have on interpretation of the wall structure. Visualisation of native, hydrated, secondary cell walls with environmental FE-SEM has been challenging and the resolution of obtained images has been low (Donaldson, 2007). We present here a technique for the analysis of native, fully-hydrated, secondary cell wall material from angiosperm and gymnosperm plant species using cryo-SEM. The use of an ultrathin 3 nm platinum film, together with cryo-preservation at high vacuum, enabled us to demonstrate that cell wall macrofibrils are a common feature in all types of native secondary cell wall material analysed. Importantly, we were able to detect the presence of macrofibrils in *Arabidopsis thaliana* vessel secondary cell walls. This allowed us to make use of the readily available cell wall-related genetic resources, revealing Arabidopsis macrofibril diameter to be dependent upon cellulose, xylan and lignin.

## Results

### Softwood and hardwood secondary cell walls contain macrofibrils

In order to investigate and compare the nanoscale architecture of gymnosperm and angiosperm cell walls we analysed stem sections taken from spruce, Ginkgo and poplar using cryo-SEM. Stems were placed in the SEM specimen stub and immediately frozen in nitrogen slush, fractured and then coated with platinum, before being passed in to the SEM chamber for imaging. Nitrogen slush is a suspension of solid nitrogen that enables high freezing rates, greatly reducing the Leidenfrost effect during plunge freezing and thus minimising structural damage (Sansinena et al., 2012). The fine grain size attributed to platinum sputtering allows small and densely packed objects to be resolved. This rapid sample preparation protocol serves to better maintain sample hydration levels and native structures for optimal EM imaging in a high vacuum environment.

We first investigated whether our cryo-SEM protocol gave comparable results to the previous FE-SEM analysis of both softwood and hardwood secondary cell walls (Donaldson, 2007). To examine if macrofibrils are found in natively hydrated, nonpretreated cell walls, cryo-SEM imaging was performed on unprocessed, frozen softwood and hardwood samples. For observing gymnosperm cell wall architecture, we first prepared softwood samples from spruce and used a low magnification to see an overview of stem cross-section (Figure 1a) and tracheid structure (Figure 1b). The inner part of the stem cross section was composed of densely packed xylem tracheids, each surrounded by cell walls. To investigate the appearance of the secondary cell walls, higher magnification images of these parts of tracheid cells were acquired. This enabled us to observe that the tracheid cell walls contain fibrous structures which frequently assembled into larger aggregates (Figure 1c and 1d, red arrows). After a further increase in magnification, individual fibrils became resolvable (Figure 1e and 1f) and their diameter was found to exceed the 3 nm diameter calculated for a single softwood elementary microfibril (Fernandes et al., 2011). Therefore the observed fibrils, if composed of cellulose, represent a higher order structure that fits the description of a “macrofibril” (Niklas, 2004, Donaldson, 2007). Similarly to spruce stem, sections from another gymnosperm, the Ginkgo, were also observed to contain macrofibrils (Figure S3). These data show that, in line with previously reported SEM imaging of dried, processed plant material (Donaldson, 2007, Terashima et al., 2009), the native, hydrated cell walls of spruce and Ginkgo also contain macrofibrils. Therefore, these structures may contribute to the higher order assembly of native gymnosperm cell walls.

**Figure 1.**
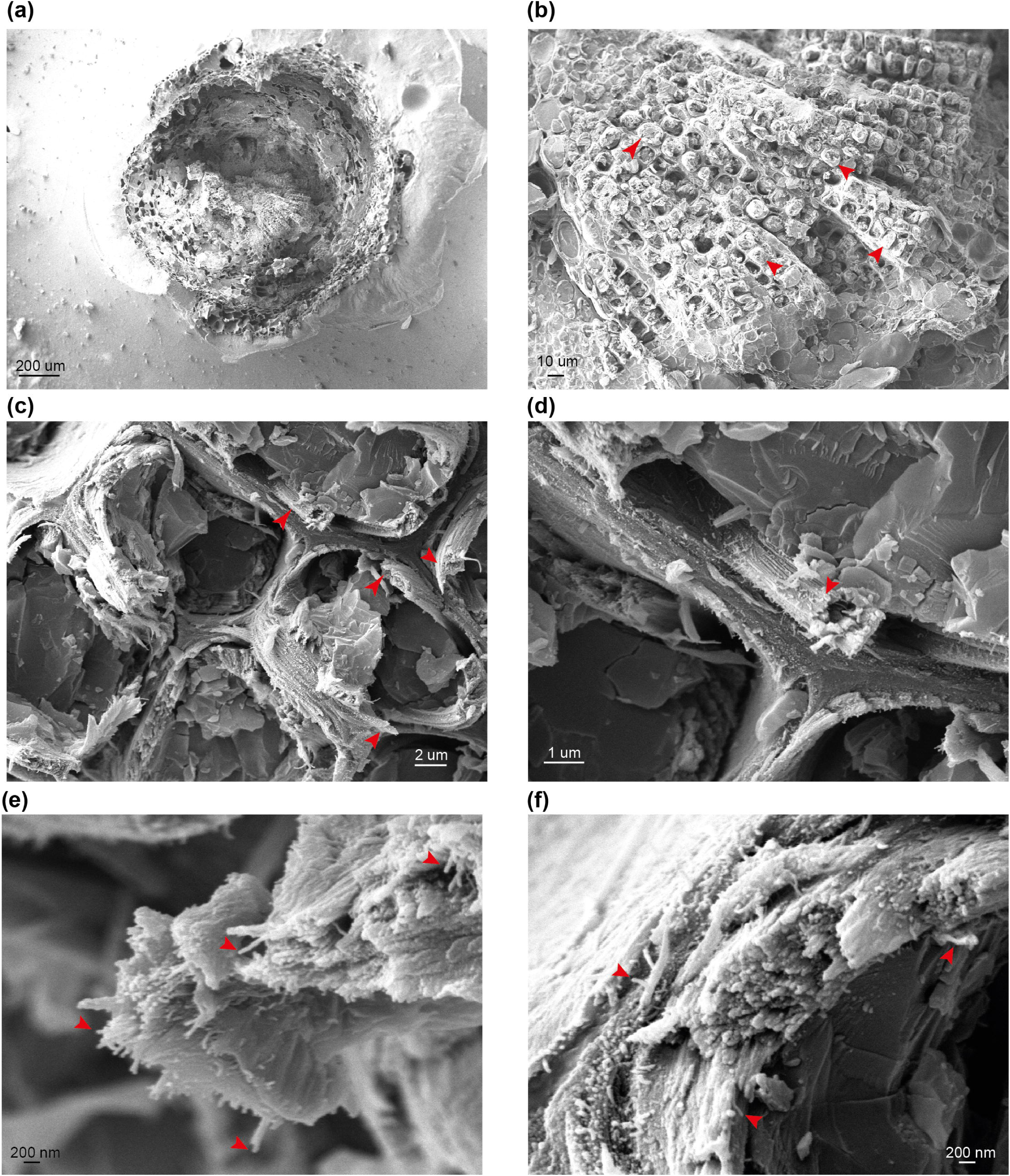
cryo-SEM analysis of spruce stem sections. **(a)** to **(f)** Representative images of stem sections of one-year-old spruce branch at different magnifications. Red arrows indicate tracheids **(b)**, macrofibril bundles **(c** and **d)** and individual macrofibrils **(e** and **f)**. Scale bars are provided for each image.

We extended the analysis to the model hardwood species, poplar. Vessels, a distinct cell type of hardwood xylem, were clearly visible using low magnification (Figure 2a and 2b). In addition to the vessels, xylem fibre cells were also observed (Figure 2b; red and yellow arrows for vessels and fibre cells respectively). For some cells we were able to observe spiral thickenings which were preserved during sample preparation and extended above the surface of the fracture plane (Figure 2b). We focused upon the vessel cell walls which showed clearly visible fibrous structures at a vessel-to-vessel boundary (Figure 2c). Analysis of vessel cell walls at a higher magnification revealed a clear presence of macrofibril structures, similar to those observed in spruce, in the poplar samples (Figure 2d and 2e). To investigate the dimensions of the macrofibrils we measured their diameter in poplar and spruce (Figure 2f). Our measurements are broadly similar to those reported in a previous study (Donaldson, 2007). We carried out comparative analysis of macrofibril diameter between hardwood and softwood by measuring 150 individual macrofibrils in poplar, spruce and Ginkgo. While the diameter of spruce and Ginkgo macrofibrils was not significantly different (Figure S4), the diameter of macrofibrils in poplar secondary cell walls was significantly smaller than that of spruce macrofibrils (Figure 2f). Spruce and Ginkgo were grown in the field while poplar samples were obtained from in vitro grown plants. To control for this difference in growth conditions we also analysed samples from field grown poplar trees. There was no statistically significant difference in the macrofibril diameter between the two poplar samples (Figure S5). For both hardwood and softwood we observed variation in the macrofibril diameter. This may reflect biological differences or may be a result of technical challenges associated with macrofibril width measurement.

**Figure 2.**
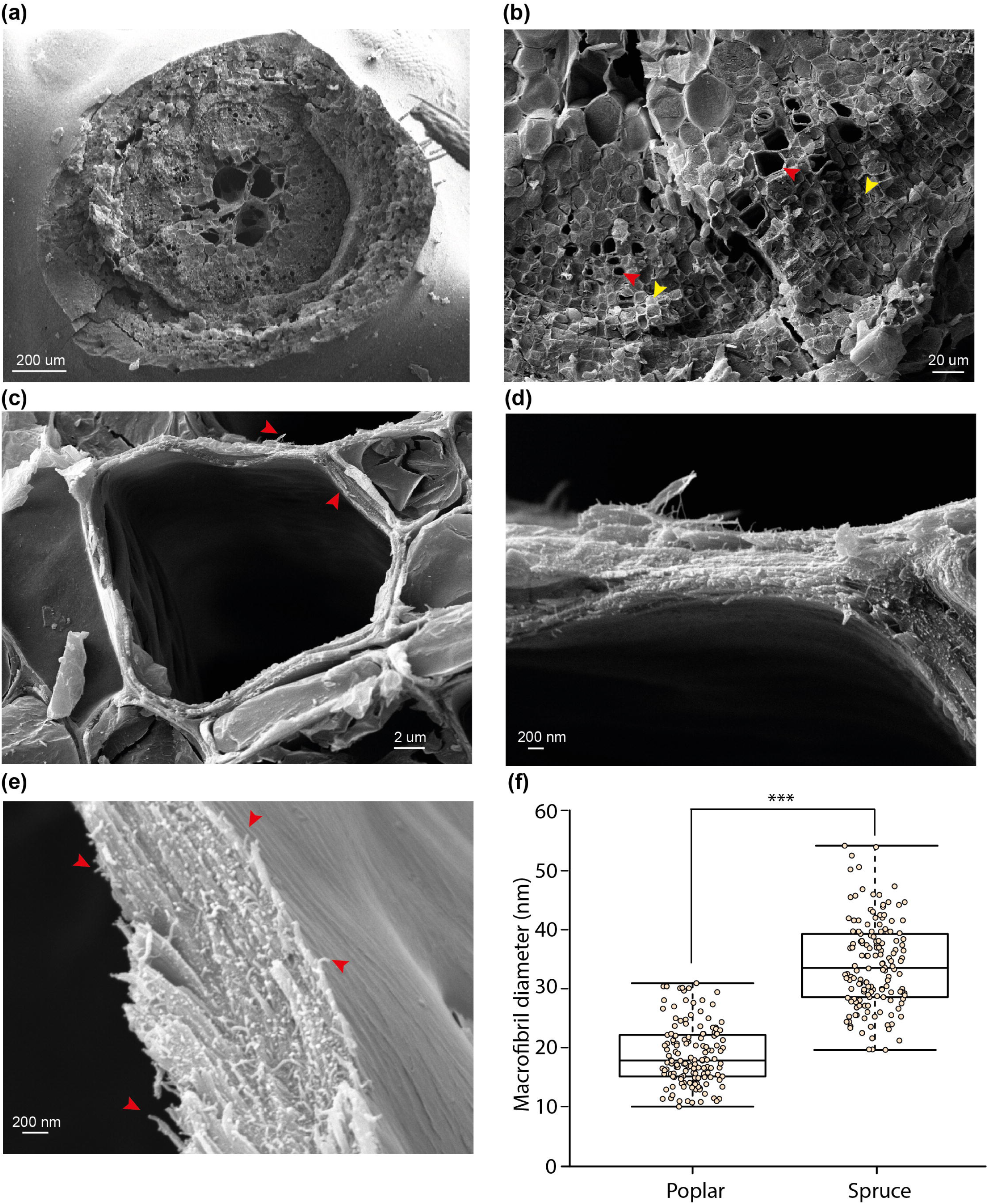
cryo-SEM analysis of poplar stem sections. **(a)** to **(e)** Representative images of stem sections of *in vitro* grown poplar trees at different magnifications. Red arrows show vessels **(b)** and macrofibrils **(c** and **e)**. Yellow arrows indicate fibre cells **(b)**. Higher magnification images **(c, d** and **e)** are presented for vessels. Scale bars are provided for each image. **(f)** Diameter of spruce tracheid cell wall fibrils compared to these observed in poplar vessel cell walls. For each bar 150 individual fibrils were measured. Boxplots mark the median and show between 25^th^ and 75^th^ percentile of the data. *** denotes p ≤ 0.00001 in Student’s t-test.

### Arabidopsis secondary cell walls macrofibrils contain a cellulose scaffold

To further evaluate the nanoscale architecture of plant cell walls and identify possible constituents of the cell wall macrofibrils, the high magnification cryo-SEM imaging was used to analyse wild type (WT) Arabidopsis secondary cell walls (Figure 3). The initial analysis investigated the structure of WT xylem vessels (Figure 3a and 3b). Sets of vessel bundles were detected and, using higher magnification, fibrous structures similar to those observed in spruce and poplar were also visible in the fractured Arabidopsis material. The width of WT Arabidopsis macrofibrils was comparable to that of poplar macrofibrils but not spruce and suggests Arabidopsis macrofibrils could be used as a good structural model for hardwoods (Figure S4, S5). Despite the use of ultra-thin platinum coating, the use of SEM without the cryo-preservation steps did not allow us to observe the Arabidopsis macrofibrils with good resolution (Figure S6) highlighting the critical importance of sample cryo-preservation to resolve a native cell wall ultrastructure.

**Figure 3.**
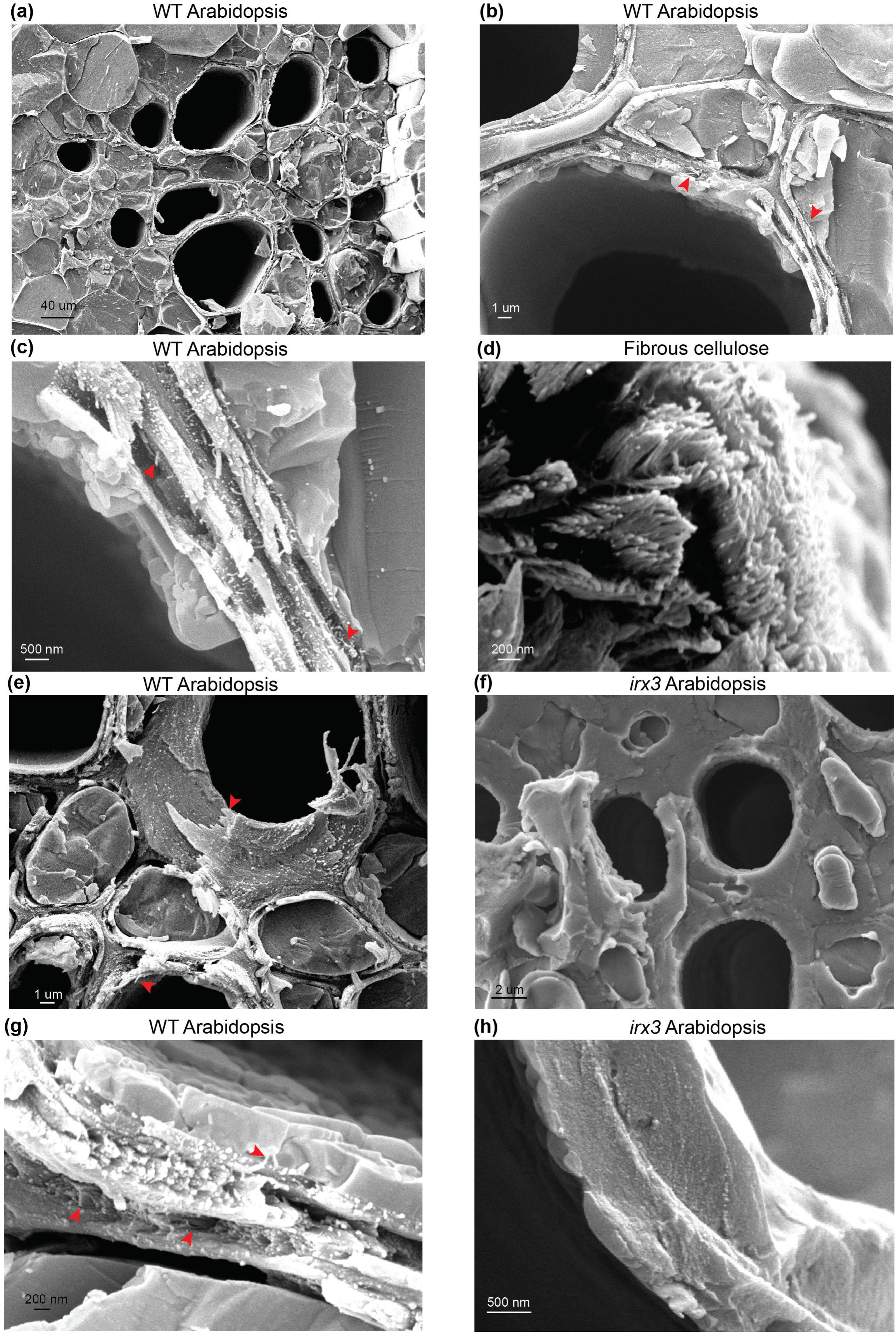
Analysis of Arabidopsis stem sections and fibrous cellulose. **(a)** to **(c)** Imaging of WT vessels at increasing magnification **(d)** Imaging of fibrous cellulose standard from cotton linters shows cell wall fibrils with an appearance similar to structures seen *in planta.* **(e)** Imaging of individual vessels in WT plants. **(f)** Imaging of individual vessels in *irx3* plants. **(g)** and **(f)** Macrofibrils are detectable in WT Arabidopsis and are absent in *irx3* secondary cell walls. Red arrows indicate the macrofibril structures throughout the figure. Scale bars are provided for each image.

Based on the data available in the literature we hypothesized that the macrofibrils may be mostly composed of cellulose (Fahlen and Salmen, 2002, Donaldson, 2007). To investigate this, and to understand the nature of these macrofibrils further, we performed a comparative analysis between WT vessel cell walls (Figure 3c) and a commercially available fibrous cellulose standard (Figure 3d) extracted from cotton linters and consisting of 99% pure cellulose (Sczostak, 2009). In this experiment, clear individual fibrils with distinct bright termini were observed in both samples indicating that the vessel wall macrofibrils have a similar appearance to the cellulose fibrils present in this polysaccharide standard. To determine whether these macrofibrils are dependent upon the proper production of cellulose, the morphology of WT Arabidopsis vessel cell walls (Figure 3e and 3g) was compared to that of the *irx3* mutant (Figure 3f and 3h). IRX3 is one of three CESA proteins that make up the secondary wall cellulose synthase complex and *irx3* plants are almost completely devoid of cellulose in their secondary cell walls, but not primary cell walls (Taylor et al., 1999). As previously reported, *irx3* plants had collapsed vessels (Figure S7), since secondary cell wall cellulose contributes to vessel wall strength (Turner & Somerville, 1997). Interestingly, the *irx3* stems lacked the fibrous structures in their vessel secondary cell walls and, in contrast to WT, the *irx3* cell walls were formed from a largely amorphous matrix (Figure 3f). It is likely that this matrix is composed of xylan and lignin, which can still be deposited in the secondary cell wall in the absence of IRX3 activity (Takenaka et al., 2018). Some structures resembling the macrofibrils were present in the primary cell walls of *irx3* plants (Figure S7). Taken together, the data demonstrate macrofibril formation is dependent upon cellulose production.

### Reduction in cell wall xylan and lignin, but not in galactoglucomannan content decreases the dimensions of Arabidopsis macrofibrils

To investigate the role of xylan in macrofibril formation, cryo-SEM was used to visualise the secondary walls from *irx9, irx10* and *esk1* Arabidopsis plants (Figure 4a, 4b and 4c). IRX9 and IRX10 are required for proper xylan synthesis and mutations in the corresponding genes lead to cell wall weakening and collapse of xylem vessels in the Arabidopsis model (Brown et al., 2007, Bauer et al., 2006, Brown et al., 2005). The *irx9* plants have impaired xylan synthesis resulting in a decrease of xylan by more than 50% compared to WT (Brown et al., 2007). In *irx10* plants the reduction in xylan content is smaller and does not exceed 20% (Brown et al., 2005). Macrofibrils are clearly observed in *irx9* and *irx10* Arabidopsis (Figure 4a and 4b). However, the median macrofibril diameter between WT and *irx9* cell wall fibres showed a ∼30% reduction in the xylan synthesis mutant (Figure 4g). The median macrofibril diameter of *irx10* plants was ∼10% smaller than that of WT Arabidopsis (Figure 4g). Although there was a wide variation in macrofibril diameter within each genotype, the difference between the WT macrofibril diameter and the one quantified for the two mutants is statistically significant, suggesting that xylan is incorporated along with cellulose to generate the normal macrofibril size. To investigate the role of xylan-cellulose interaction in the macrofibril formation we assessed the macrofibril size in the *esk1* Arabidopsis mutant (Figure 4c). Mutation in the *ESK1* gene results in reduction of xylan acetylation, but not in a decrease in xylan quantity (Xiong et al., 2013), which leads to changes in xylan [Me]GlcA patterning and loss of interaction between xylan and the hydrophilic surface of the cellulose microfibril (Grantham et al., 2017). In line with the results observed for *irx9* and *irx10* plants the loss of xylan-cellulose interaction caused a reduction in the macrofibril diameter (Figure 4g).

**Figure 4.**
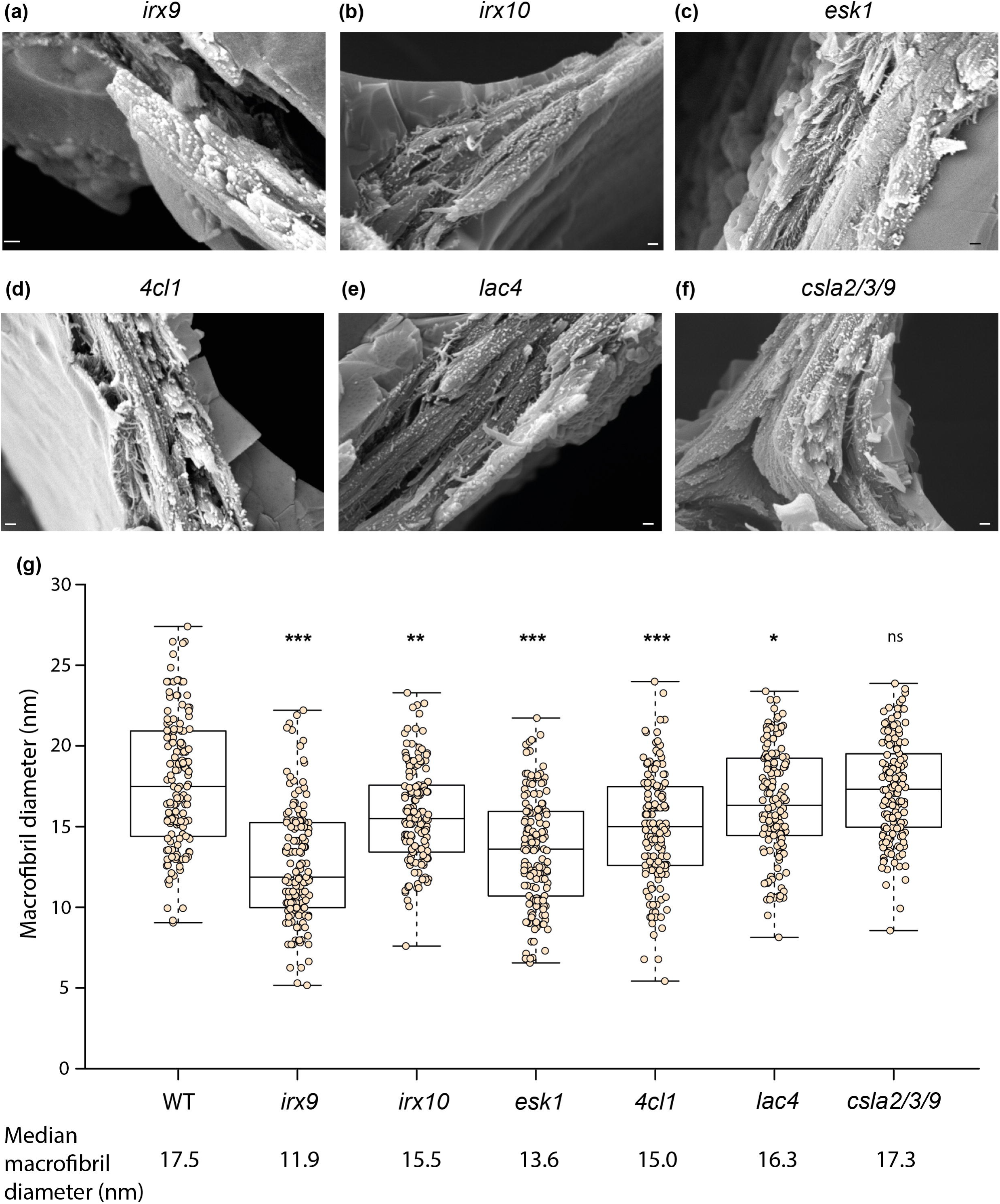
Analysis of macrofibrils in mutant Arabidopsis plants. Representative image of **(a)** *irx9*, **(b)** *irx10*, **(c)** *esk1*, **(d)** *4cl1*, **(e)** *lac4* and **(f)** *csla2/3/9* Arabidopsis macrofibrils. Scale bar corresponds to 200 nm on each image. **(g)** Quantification of macrofibril diameter in WT and mutant Arabidopsis plants. N = 150. Boxplots mark a median and show between 25^th^ and 75^th^ percentile of the data. *** denotes p ≤ 0.00001, ** denotes p ≤ 0.0001, * denotes p ≤ 0.05 in Tukey test following ANOVA when compared to WT, ns indicates lack of statistically significant difference.

Previous work in softwood suggested that lignin (Donaldson, 2007) and galactoglucomannan (GGM) (Terashima et al., 2009) may be involved in macrofibril formation. To investigate the role of these two cell wall components in the maintenance of macrofibril structure we performed imaging of *4cl1* (Figure 4d), *lac4* (Figure 4e) and *csla2/3/9* (Figure 4f) mutant Arabidopsis cell walls. Both 4CL1 and LAC4 are involved in lignin biosynthesis and plants mutated in genes encoding these enzymes have a 30% and 15% reduction in lignin content respectively (Li et al., 2015, Berthet et al., 2011). The median macrofibril diameter for both *4cl1* and *lac4* was significantly smaller than that calculated for WT (Figure 4g). Importantly, the extent of the reduction in macrofibril diameter was in line with the decrease in the lignin content observed for the two mutants, with *4cl1* macrofibrils being ∼15% smaller than the WT ones and *lac4* macrofibrils having ∼7% reduction in the median diameter. Proteins from the CSLA family are involved in the biosynthesis of a hemicellulose galactoglucomannan and mutations in *csla2/3/9* leads to nearly complete loss of stem GGM in the Arabidopsis model (Goubet et al., 2009). Our quantitative analysis indicates that the diameter of macrofibrils of *csla2/3/9* Arabidopsis was not significantly different to that of the WT plants (Figure 4g).

## Discussion

The native nanoscale architecture of woody plant secondary cell walls remains poorly understood due to the challenges of keeping the sample hydrated, which is incompatible with some types of techniques. Studies analysing dehydrated and fixed plant cell wall samples with FE-SEM (Donaldson, 2007), together with other work which includes SANS experiments investigating spruce (Fernandes et al., 2011) and bamboo samples (Thomas et al., 2015), suggest there is a higher order arrangement of cellulose microfibrils in plant secondary cell walls. Our work reports the application of a cryo-SEM based analysis technique which, using exclusively samples that have not been dried, heated or chemically processed, indicates that secondary cell wall cellulose microfibrils are likely to come together to form larger macrofibril structures.

### Our study strongly suggests that these structures, at least in the model plant species *Arabidopsis thaliana*, contain cellulose, xylan and lignin

Previous studies investigated the presence and diameter of macrofibrils in dehydrated softwood samples (Donaldson, 2007). In line with results presented in our work, Donaldson did observe macrofibrils in cell walls of pine tracheids. Moreover, also in agreement with the results presented here (Figure S4), these softwood macrofibrils were larger than those seen in hardwoods. In softwood, in addition to various patterned types of xylan (Busse-Wicher et al., 2016b, Martinez-Abad et al., 2017), most of which are likely to be compatible with binding to the hydrophilic surface of the cellulose fibril, the cell walls contain large quantities of acetylated GGM (Scheller and Ulvskov, 2010) which may contribute to macrofibril width. Indeed, gymnosperm GGM was proposed to interact with the cellulose microfibril in cell walls of Ginkgo (Terashima et al., 2009). Therefore, the significant difference in macrofibril diameter observed between hardwood and softwood samples may be due to the differences in the cell wall composition. Consequently, we hypothesize that in gymnosperms, GGM, along with xylan, may contribute to the macrofibril size in a way similar to what we observed for xylan in Arabidopsis macrofibrils. With an average diameter ranging between 20 and 34 nm, the size of pine macrofibrils measured by Donaldson was somewhat smaller than that measured in spruce wood in the current work. However, these observations are not necessarily inconsistent. Donaldson dehydrated the wood samples prior to the SEM imaging. As the spacing between bundled softwood cellulose microfibrils, estimated to be equal to 3 nm by small angle neutron scattering, is sensitive to wood hydration levels (Fernandes et al., 2011), at least part of the difference in the macrofibril diameter might be due to the changes in the water content within the sample analysed with SEM. Interestingly, Donaldson reported that macrofibrils in dried poplar wood, depending on their position in cell wall, have an average diameter ranging from 14 to 18 nm, which is similar to what was measured for both poplar and Arabidopsis as a part of our study. This observation suggests that the softwood macrofibril size may be more sensitive to drying than the hardwood one. This in turn suggests that, in addition to compositional disparities, hydration could contribute to the differences in softwood and hardwood macrofibril characteristics. In addition to providing scientific insight, this result highlights that imaging of the cryopreserved secondary cell walls offers significant advance over the previously used techniques.

Interestingly, similar to a previous report (Donaldson, 2007), we observed that macrofibrils in both hardwood and softwood have a range of diameters. The reasons for this variation in size are not clear. It is possible that the number of individual cellulose microfibrils that come together to form the macrofibril structure in both hardwood and softwood is not constant. This may be regulated by coordinated movement of CesA complexes or their density during cell wall synthesis (Li et al., 2016). It was proposed that the macrofibril diameter is proportional to the degree of cell wall lignification (Donaldson, 2007), which may also vary between the structures. This hypothesis is supported by our results which indicate that the cell wall lignin content influences macrofibril diameter in Arabidopsis. Variations may also originate from environmental conditions. For example, it was shown that wood density may vary correlatively with climate change (Bouriaud et al., 2005). Although much of this effect is likely to be due to cell size and wall thickness, it can be hypothesized that change in wood density may also originate from compositional changes that impact macrofibril assembly and ultrastructure. It would therefore be relevant to assess macrofibrils of perennial trees with samples spanning several years of growth. We cannot rule out that the width variance may originate from the technical limitations of resolving the macrofibrils by SEM. It will be interesting to see if the emerging He-ion technologies, with an increase in resolution and less dependent upon metal coating, reduce this variance (Joens et al., 2013). The cryo-SEM techniques developed as part of our study offer a significant advantage over the previous investigation (Donaldson, 2007) which applied a thicker coat of chromium (mostly 12 nm) that yield films with coarser grains than the thinner (3 nm) platinum films used in our work. Thus, taking the results described by Donaldson and our technological improvements into consideration, we believe that the variance in the macrofibril width observed in both studies is likely to reflect natural material variation.

The prominence of macrofibril structures in Arabidopsis cell walls is a surprising discovery of this study. Previously published results using AFM analysis indicate the presence of some bundled microfibrils in primary cell walls of Arabidopsis but the extent of this bundling is lower than what was observed in primary cell wall samples from other species (Zhang et al., 2016). AFM is not yet technically feasible for analysis of bundling of hydrated secondary cell walls although recent technical advances allowed visualisation of dried spruce wood at a nanometer resolution (Casdorff et al., 2017). The observation of the macrofibrils by cryo-SEM in Arabidopsis allowed us to determine the contribution of cellulose, xylan, lignin and galactoglucomannan to macrofibril formation, thanks to the availability of secondary cell wall related mutants in this model. Macrofibrils were completely absent in vessel cell walls of *irx3* plants, which lack secondary cell wall cellulose, indicating that proper cellulose biosynthesis is required for formation and assembly of secondary cell walls polymers into macrofibrils. In addition, we observed that vessel macrofibril diameter is significantly decreased in *irx9, irx10* and *esk1* plants, suggesting that xylan may also participate in the correct assembly of such structures. While in *irx9* and *irx10* reduction in macrofibril diameter may be associated with decrease in the xylan content the ∼25% reduction in the median macrofibril diameter observed for *esk1* Arabidopsis is harder to explain. Hardwood xylan is proposed to interact with the hydrophilic surface of the cellulose microfibril as a two-fold screw (Simmons et al., 2016, Busse-Wicher et al., 2016a), and this interaction is facilitated by the even pattern of the [Me]GlcA and acetyl branches on the xylan backbone which is lost in *esk1* plants (Grantham et al., 2017). Therefore, the decrease in macrofibril diameter observed in *esk1* Arabidopsis indicates that xylancellulose interaction may have a role in spacing or proper coalescence of microfibrils to form the elementary macrofibril. It is unclear why the macrofibril diameter is reduced in *esk1*, but perhaps fewer elementary fibrils are incorporated into each macrofibril when xylan is not interacting with the hydrophilic surface of the cellulose fibril. This may be different to the effect observed in flax where the absence of xylan may lead to aggregation of glucan chains into larger fibres (Thomas et al., 2013). Such difference may be associated with variations in the stoichiometry of the cellulose synthase complex which were recently reported for angiosperms (Zhang et al., 2018).

In addition to implicating xylan in the process of macrofibril formation our results indicate that lignin may contribute to assembly of the structures. As such, our results use genetic assignment to extend previous work which has correlated macrofibril diameter with the degree of wall lignification (Donaldson, 2007). Interestingly, we observed that the macrofibril diameter does not correlate with the cell wall GGM content. This may be associated with low abundance of GGM in angiosperms where the polysaccharide accounts for only up to 5% of the cell wall material (Scheller and Ulvskov, 2010). Alternatively, this result may indicate that in Arabidopsis GGM might be not involved in macrofibril formation. GGM may play a more significant role in the macrofibril assembly in gymnosperms where it accounts for up to 30% of the cell wall material. Importantly, all our conclusions are based on the analysis of native, hydrated, cell wall samples. The assignment of cell wall macrofibril composition, in their native state, would be impossible using techniques such as immunogold due to the pretreatment steps needed before the antibody labelling.

In conclusion, our analysis indicates that vessel cell walls contain fibrous structures composed of cellulose, xylan and lignin. These structures are present in both hardwood and softwood and have a diameter larger than a single cellulose microfibril. Therefore, these structures can be described as cell wall macrofibrils. The reduction in macrofibril diameter observed in *esk1* Arabidopsis suggests that the interaction between xylan and the hydrophilic surface of the cellulose microfibril may be involved in the assembly of these structures. Therefore, this xylan-cellulose interaction may be important for the maintenance of plant cell wall ultrastructure and mechanical properties (Simmons et al., 2016). The techniques developed here and the discovery of the ubiquitous presence of macrofibrils in hardwood and softwood in their native state will contribute to a better understanding of cell wall assembly processes. Furthermore, the ability to resolve macrofibrils in Arabidopsis, along with the availability of genetic resources in this model, will offer the community a valuable tool to further study the complex deposition of secondary cell walls polymers and their role in defining the cell wall ultrastructure. The assembly of cell wall macrofibrils is likely to influence the properties of wood, such as density, which may vary due to different stimuli such as tree fertilisation (Makinen et al., 2002) or environmental changes (Bouriaud et al., 2005). Therefore, we expect that the methodology described here will enable to correlate the native nanoscale features of the cell walls, such as the macrofibril diameter, or a specific macrofibril patterning within the cell wall, with wood properties. Consequently, our approach may be useful to assess this aspect of wood quality at a new level and could benefit numerous industries ranging from building construction, paper manufacturing and biofuel production to generation of novel biomaterials such as nanocrystalline cellulose.

## Experimental procedures

### Plant material

*Picea abies*, (spruce) one-year old branch was acquired from 30-50cm tall potted plants grown outdoors purchased from Scotsdale (Great Shelford, Cambridgeshire, UK). *Ginkgo biloba*, (Ginkgo) stem material was obtained from the trees grown at the Cambridge University Botanic Garden. For both spruce and Ginkgo, samples from two individuals were analysed.

Hybrid aspen (*Populus tremula* × *Populus tremuloides*, clone T89), referred to as poplar in the text, was grown *in vitro* (20°C, with a 16-h light, 8-h dark photoperiod, with illumination at 85 microeinstein.m^-2^.s^-1^) during 76 to 80 days after micropropagation on 1/2MS media with vitamins (Duchefa M0222), 1% sucrose, 0.7% Agar. Samples from three individuals were analysed. For field grown poplar (*Populus tremula*), material was obtained from two individuals grown at the Cambridge University Botanic Garden.

*Arabidopsis thaliana* (Arabidopsis) Columbia-0 ecotype plants were grown in a cabinet maintained at 21 °C, with a 16-h light, 8-h dark photoperiod. Stem material was collected from 7-week-old plants. Mutant insertion lines described in published work were used in this study. Specifically, *irx3-7* plants (Simmons et al., 2016, Kumar and Turner, 2015), representing a mutant allele of *CESA7, irx9-1* (Brown et al., 2005), *irx10-1* (Brown et al., 2005), *esk1-5* (Lefebvre et al., 2011, Grantham et al., 2017), *4cl1-1* (Vanholme et al., 2012), *lac4-2* (Berthet et al., 2011) and *csla2-1csla3-2csla91* (Goubet et al., 2009). Plants were analysed alongside the wild type (WT) material. For each genotype three individuals were analysed.

### Cryo-SEM sample preparation and imaging

Fresh stems of 7 week old Arabidopsis plants were prepared for imaging as outlined in Supporting Information Figure S1. Firstly, 1 cm length sections were cut from the bottom part of the stems and mounted vertically in recessed stubs containing a cryo glue preparation consisting of a 3:1 mixture of Tissue-Tec (Scigen Scientific, USA) and Aquadog colloidal graphite (Agar Scientific, Stansted, UK) (see steps 1 to 4 on Figure S1). Stem sections were immediately (within 5 minutes of harvest) plunge frozen in liquid nitrogen slush (step 5 on Figure S1), transferred under vacuum, fractured and then coated with 3 nm of platinum (step 6 on Figure S1) using a PT3010T cryo-apparatus fitted with a film thickness monitor (Quorum Technologies, Lewes, UK). The short time between freezing and harvesting serves to prevent drying of the sample where only the exposed surface, not the fractured face, is expected to exhibit some water loss during the short time it is exposed to air. Finally, fractured stems were imaged using a Zeiss EVO HD15 Scanning Electron Microscope (step 7 on Figure S1) and maintained at −145 °C using a Quorum cryo-stage assembly. The electron source is a Lanthanum Hexaboride HD filament. Images were acquired using a secondary electron detector and an accelerating voltage of between 5 and 8 kV with a working distance between 4 and 6 mm. Quantification of the width of cell wall macrofibrils was performed using ImageJ software (Schneider et al., 2012). For the measurements of each macrofibril, a line was drawn parallel to the fibril axis. The length of a second line, perpendicular to the fibril axis line and across the width of the macrofibril, was quantified as the macrofibril width (Figure S2). Each fibril width measurement was standardised for the platinum layer applied during the coating process by subtracting the width of the standardised coat from the original measurement. Imaging without the cryo-preservation was performed by visualising hand sectioned platinum coated specimens with the stage maintained at room temperature. For preparation of these samples all freezing steps were omitted.

### Sampling and statistical analysis

For spruce, Ginkgo and field grown poplar, stem sections were taken from two individual trees and 150 macrofibrils were measured from three tracheids that had each been coated with platinum separately. Imaging of poplar was performed in technical triplicate from three *in vitro* grown trees and 150 poplar macrofibrils were measured from three separately coated vessels as for the gymnosperm samples. For Arabidopsis, cryo-SEM imaging of vessels was carried out on three biological replicates, each from separate individuals. 150 macrofibril diameters were measured across the three individuals.

Statistical analysis was performed using packages available with R software (Team, 2014). Statistical tests, either Student’s T test or ANOVA, used to compare average measurements for samples are defined in Figure legends. The variance between each pairwise combination was estimated to be similar with Levene’s test.

### Data statement

All data for the quantification of the macrofibril diameter is presented on Figures forming part of the manuscript. Representative images are provided for each genotype/species for which macrofibril diameter data is provided. All images obtained for the different species and genotypes analysed as part of this manuscript are available from Jan J Lyczakowski.

## Accession numbers

Mutants in the following genes in the Col-0 ecotype were analysed as part of this study:

AT5G17420 (*irx3-7*)

AT2G37090 (*irx9-1*)

AT1G27440 (*irx10-1*)

AT3G55990 (*esk1-5*)

AT1G51680 (*4cl1-1*)

AT2G38080 (*lac4-2*)

AT5G22740 (*csla2-1*)

AT1G23480 (*csla3-2*) AT5G03760

(*csla9-1*)

## List of abbreviations

1D: one dimensional
AFM: atomic force microscopy CesA – Cellulose
synthase cryo-SEM: low temperature scanning electron
microscopy FE-SEM: field emission scanning electron microscopy
FT-IR: Fourier-transform infrared spectroscopy
GGM: galactoglucomannan
He-ion: Helium ion
IRX: irregular xylem
[Me]GlcA: methylated and unmethylated form of glucuronic acid
NMR: nuclear magnetic resonance
SANS: small angle neutron scattering
TEM: transmission electron microscopy
WAXS: wide angle x-ray scattering

## Acknowledgements

We would like to thank Dr Nadine Anders and Dr Henry Temple for helpful discussions and critical reading of the manuscript. This work was supported by the Leverhulme Trust Centre for Natural Material Innovation. Analysis of Arabidopsis mutant plants was supported as part of The Center for LignoCellulose Structure and Formation, an Energy Frontier Research Center funded by the U.S. Department of Energy (DOE), Office of Science, Basic Energy Sciences (BES), under Award # DE-SC0001090. JJL was in receipt of a studentship from the Biotechnology and Biological Sciences Research Council (BBSRC) of the UK as part of the Cambridge BBSRC-DTP Programme (Reference BB/J014540/1). MB is employed in YH’s team through the European Research Council Advanced Investigator Grant SYMDEV (No. 323052). YH laboratory is funded by the Finnish Centre of Excellence in Molecular Biology of Primary Producers (Academy of Finland CoE program 2014-2019) (decision #271832); the Gatsby Foundation (GAT3395/PR3); the National Science Foundation Biotechnology and Biological Sciences Research Council grant (BB/N013158/1); University of Helsinki (award 799992091), and the European Research Council Advanced Investigator Grant SYMDEV (No. 323052). OMT was a recipient of an iCASE studentship from the BBSRC (Reference BB/M015432/1). We thank Gareth Evans for help with cryo-SEM sample preparation. The cryo-SEM facility at the Sainsbury Laboratory is supported by the Gatsby Charitable Foundation. We are grateful to Cambridge University Botanic Garden for access to the *Populus tremula* and *Ginkgo biloba* collections.

The authors declare no conflict of interest.

## Author contributions

JJL designed the study, conducted the experiments, analysed the data and wrote the paper. MB performed poplar imaging experiments, analysed the data and wrote the paper. OMT analysed the data and wrote the paper. YH contributed to data analysis and manuscript preparation, RW designed the study, conducted experiments, analysed the data and wrote the paper. PD designed the study and contributed to data analysis and manuscript preparation.

## Short legends for supporting information

Figure S1: Overview of the cryo-SEM procedure.

Figure S2: Measurement of macrofibril diameter.

Figure S3: Cryo-SEM analysis of Ginkgo cell walls.

Figure S4: Comparison of macrofibril diameter in Arabidopsis, poplar, spruce and Ginkgo.

Figure S5: Imaging of macrofibrils in field grown poplar.

Figure S6: Analysis of native Arabidopsis samples without the cryo-preservation protocol.

Figure S7: Cryo-SEM analysis of vessel collapse and primary cell wall cellulose in *irx3* Arabidopsis plants.

## References

Abe, H., Funada, R., Ohtani, J. & Fukazawa, K. 1997. Changes in the arrangement of cellulose microfibrils associated with the cessation of cell expansion in tracheids. Trees-Structure and Function, 11, 328–332.

Bauer, S., Vasu, P., Persson, S., Mort, A. J. & Somerville, C. R. 2006. Development and application of a suite of polysaccharide-degrading enzymes for analyzing plant cell walls. Proceedings of the National Academy of Sciences of the United States of America, 103, 11417–11422.

Berthet, S., Demont-Caulet, N., Pollet, B., Bidzinski, P., Cezard, L., Le Bris, P., Borrega, N., Herve, J., Blondet, E., Balzergue, S., Lapierre, C. & Jouanin, L. 2011. Disruption of LACCASE4 and 17 Results in Tissue-Specific Alterations to Lignification of Arabidopsis thaliana Stems. Plant Cell, 23, 1124–1137.

Bonawitz, N. D. & Chapple, C. 2010. The Genetics of Lignin Biosynthesis: Connecting Genotype to Phenotype. Annual Review of Genetics, Vol 44, 44, 337–363.

Bouriaud, O., Leban, J. M., Bert, D. & Deleuze, C. 2005. Intra-annual variations in climate influence growth and wood density of Norway spruce. Tree Physiology, 25, 651–660.

Bromley, J. R., Busse-Wicher, M., Tryfona, T., Mortimer, J. C., Zhang, Z., Brown, D. M. & Dupree, P. 2013. GUX1 and GUX2 glucuronyltransferases decorate distinct domains of glucuronoxylan with different substitution patterns. Plant Journal, 74, 423–434.

Brown, D. M., Goubet, F., Vicky, W. W. A., Goodacre, R., Stephens, E., Dupree, P. & Turner, S. R. 2007. Comparison of five xylan synthesis mutants reveals new insight into the mechanisms of xylan synthesis. Plant Journal, 52, 1154–1168.

Brown, D. M., Zeef, L. A. H., Ellis, J., Goodacre, R. & Turner, S. R. 2005. Identification of novel genes in Arabidopsis involved in secondary cell wall formation using expression profiling and reverse genetics. Plant Cell, 17, 2281–2295.

Busse-Wicher, M., Gomes, T. C. F., Tryfona, T., Nikolovski, N., Stott, K., Grantham, N. J., Bolam, D. N., Skaf, M. S. & Dupree, P. 2014. The pattern of xylan acetylation suggests xylan may interact with cellulose microfibrils as a twofold helical screw in the secondary plant cell wall of Arabidopsis thaliana. Plant Journal, 79, 492–506.

Busse-Wicher, M., Grantham, N. J., Lyczakowski, J. J., Nikolovski, N. & Dupree, P. 2016a. Xylan decoration patterns and the plant secondary cell wall molecular architecture. Biochemical Society Transactions, 44, 74–78.

Busse-Wicher, M., Li, A., Silveira, R. L., Pereira, C. S., Tryfona, T., Gomes, T. C. F., Skaf, M. S. & Dupree, P. 2016b. Evolution of Xylan Substitution Patterns in Gymnosperms and Angiosperms: Implications for Xylan Interaction with Cellulose. Plant Physiology, 171, 2418–2431.

Casdorff, K., Keplinger, T., Burgert, I. 2017. Nono-mechanical characterisation of the wood cell wall by AFM studies: comparison between AC- and QITM mode. Plant Methods, 13, 60.

Cochard, H., Froux, F., Mayr, F. F. S. & Coutand, C. 2004. Xylem wall collapse in water-stressed pine needles. Plant Physiology, 134, 401–408.

Cosgrove, D. J. 2014. Re-constructing our models of cellulose and primary cell wall assembly. Current Opinion in Plant Biology, 22, 121–131.

Dick-Perez, M., Zhang, Y. A., Hayes, J., Salazar, A., Zabotina, O. A., Hong, M. 2011. Structure and interactions of plant cell-wall polysaccharides by two- and three-dimensional magic-angle-spinning solid-state NMR. Biochemistry, 50, 989–1000.

Donaldson, L. 2007. Cellulose microfibril aggregates and their size variation with cell wall type. Wood Science and Technology, 41, 443–460.

Elbaum, R., Gorb, S., Fratzl, P. 2008. Structures in the cell wall that enable hygroscopic movement of wheat awns. Journal of Structural Biology, 164, 101–107.

Fahlen, J. & Salmen, L. 2002. On the Lamellar Structure of the Tracheid Cell Wall. Plant Biology, 4, 339–345.

Favery, B., Ryan, E., Foreman, J., Linstead, P., Boudonck, K., Steer, M., Shaw, P. & Dolan, L. 2001. KOJAK encodes a cellulose synthase-like protein required for root hair cell morphogenesis in Arabidopsis. Genes & Development, 15, 79–89.

Fernandes, A. N., Thomas, L. H., Altaner, C. M., Callow, P., Forsyth, V. T., Apperley, D. C., Kennedy, C. J. & Jarvis, M. C. 2011. Nanostructure of cellulose microfibrils in spruce wood. Proceedings of the National Academy of Sciences of the United States of America, 108, E1195E1203.

Fromm, J., Rockel, B., Lautner, S., Windeisen, E. & Wanner, G. 2003. Lignin distribution in wood cell walls determined by TEM and backscattered SEM techniques. Journal of Structural Biology, 143, 77–84.

Fujita, M., Himmelspach, R., Ward, J., Whittington, A., Hasenbein, N., Liu, C., Truong, T. T., Galway, M. E., Mansfield, S. D., Hocart, C. H. & Wasteneys, G. O. 2013. The anisotropy1 D604N Mutation in the Arabidopsis Cellulose Synthase1 Catalytic Domain Reduces Cell Wall Crystallinity and the Velocity of Cellulose Synthase Complexes. Plant Physiology, 162, 74–85.

Gierlinger, N. 2018. New insights into plant cell walls by vibrational microspectroscopy. Applied Spectroscopy Reviews, 53, 517–551.

Gonneau, M., Desprez, T., Guillot, A., Vernhettes, S. & Hofte, H. 2014. Catalytic Subunit Stoichiometry within the Cellulose Synthase Complex. Plant Physiology, 166, 1709–1712.

Goubet, F., Barton, C. J., Mortimer, J. C., Yu, X. L., Zhang, Z. N., Miles, G. P., Richens, J., Liepman, A. H., Seffen, K. & Dupree, P. 2009. Cell wall glucomannan in Arabidopsis is synthesised by CSLA glycosyltransferases, and influences the progression of embryogenesis. Plant Journal, 60, 527–538.

Grantham, N. J., Wurman-Rodrich, J., Terrett, O. M., Lyczakowski, J. J., Stott, K., Iuga, D., Simmons, T. J., Durand-Tardif, M., Brown, S. P., Dupree, R., Busse-Wicher, M. & Dupree, P. 2017. An even pattern of xylan substitution is critical for interaction with cellulose in plant cell walls. Nature Plants, 3, 859–865.

Hill, J. L., Hammudi, M. B. & Tien, M. 2014. The Arabidopsis Cellulose Synthase Complex: A Proposed Hexamer of CESA Trimers in an Equimolar Stoichiometry. Plant Cell, 26, 4834–4842.

Himmelspach, R., Williamson, R. E. & Wasteneys, G. O. 2003. Cellulose microfibril alignment recovers from DCB-induced disruption despite microtubule disorganization. Plant Journal, 36, 565–575.

Jarvis, M. C. 2018. Structure of native cellulose microfibrils, the starting point for nanocellulose manufacture. Philosophical transactions. Series A, Mathematical, physical, and engineering sciences, 376.

Joens, M. S., Huynh, C., Kasuboski, J. M., Ferranti, D., Sigal, Y. J., Zeitvogel, F., Obst, M., Burkhardt, C. J., Curran, K. P., Chalasani, S. H., Stern, L. A., Goetze, B. & Fitzpatrick, J. A. J. 2013. Helium Ion Microscopy (HIM) for the imaging of biological samples at sub-nanometer resolution. Scientific Reports, 3.

Kang, X., Kirui, A., Dickwella Widanage, M. C., Mentink-Vigier, F., Cosgrove, D. J., Wang, T. 2019. Lignin-polysaccharide interactions in plant secondary cell walls revealed by solid-state NMR. Nature Communications, 10, 347.

Kumar, M. & Turner, S. 2015. Protocol: a medium-throughput method for determination of cellulose content from single stem pieces of Arabidopsis thaliana. Plant Methods, 11.

Lefebvre, V., Fortabat, M. N., Ducamp, A., North, H. M., Maiagrondard, A., Trouverie, J., Boursiac, Y., Mouille, G. & Durand-Tardif, M. 2011. ESKIMO1 Disruption in Arabidopsis Alters Vascular Tissue and Impairs Water Transport. Plos One, 6.

Li, S. D., Bashline, L., Zheng, Y. Z., Xin, X. R., Huang, S. X., Kong, Z. S., Kim, S. H., Cosgrove, D. J. & Gu, Y. 2016. Cellulose synthase complexes act in a concerted fashion to synthesize highly aggregated cellulose in secondary cell walls of plants. Proceedings of the National Academy of Sciences of the United States of America, 113, 11348–11353.

Li, Y., Kim, J. I., Pysh, L. & Chapple, C. 2015. Four Isoforms of Arabidopsis 4Coumarate: CoA Ligase Have Overlapping yet Distinct Roles in Phenylpropanoid Metabolism. Plant Physiology, 169, 2409–2421.

Loque, D., Scheller, H. V. & Pauly, M. 2015. Engineering of plant cell walls for enhanced biofuel production. Current Opinion in Plant Biology, 25, 151–161.

Lyczakowski, J. J., Wicher, K. B., Terrett, O. M., Faria-Blanc, N., Yu, X. L., Brown, D., Krogh, K., Dupree, P. & Busse-Wicher, M. 2017. Removal of glucuronic acid from xylan is a strategy to improve the conversion of plant biomass to sugars for bioenergy. Biotechnology for Biofuels, 10.

Makinen, H., Saranpaa, P. & Linder, S. 2002. Wood-density variation of Norway spruce in relation to nutrient optimization and fibre dimensions. Canadian Journal of Forest Research-Revue Canadienne De Recherche Forestiere, 32, 185–194.

Marga, F., Grandbois, M., Cosgrove, D. J. & Baskin, T. I. 2005. Cell wall extension results in the coordinate separation of parallel microfibrils: evidence from scanning electron microscopy and atomic force microscopy. Plant Journal, 43, 181–190.

Martinez-Abad, A., Berglund, J., Toriz, G., Gatenholm, P., Henriksson, G., Lindstrom, M., Wohlert, J. & Vilaplana, F. 2017. Regular Motifs in Xylan Modulate Molecular Flexibility and Interactions with Cellulose Surfaces. Plant Physiology, 175, 1579–1592.

Niklas, K. 2004. The cell walls that bind the tree of life. BioScience, 54, 831-841.

Nishimura, H., Kamiya, A., Nagata, T., Katahira, M. & Watanabe, T. 2018. Direct evidence for alpha ether linkage between lignin and carbohydrates in wood cell walls. Scientific Reports, 8.

Pan, Y. D., Birdsey, R. A., Fang, J. Y., Houghton, R., Kauppi, P. E., Kurz, W. A., Phillips, O. L., Shvidenko, A., Lewis, S. L., Canadell, J. G., Ciais, P., Jackson, R. B., Pacala, S. W., Mcguire, A. D., Piao, S. L., Rautiainen, A., Sitch, S. & Hayes, D. 2011. A Large and Persistent Carbon Sink in the World’s Forests. Science, 333, 988–993.

Pauly, M. & Keegstra, K. 2008. Cell-wall carbohydrates and their modification as a resource for biofuels. Plant Journal, 54, 559–568.

Purbasha, S., Bosneaga, E. Auer, M. 2009. Plant cell walls throughout evolution: towards a molecular understanding of their design principles. Journal of Experimental Botany, 60, 3615–3635.

Ralph, J., Lundquist, K., Brunow, G., Lu, F., Kim, H., Schatz, P. F., Marita, J. M., Hatfield, R. D., Ralph, S. A., Christensen, J. H., Boerjan, W. 2004. Lignins: natural polymers from oxidative coupling of 4hydroxyphenyl-propanoids. Phytochemistry Reviews, 3, 29–60.

Ramage, M. H., Burridge, H., Busse-Wicher, M., Fereday, G., Reynolds, T., Shah, D. U., Wu, G. L., Yu, L., Fleming, P., Densleytingley, D., Allwood, J., Dupree, P., Linden, P. F. & Scherman, O. 2017. The wood from the trees: The use of timber in construction. Renewable & Sustainable Energy Reviews, 68, 333–359.

Refregier, G., Pelletier, S., Jaillard, D. & Hofte, H. 2004. Interaction between wall deposition and cell elongation in dark-grown hypocotyl cells in Arabidopsis. Plant Physiology, 135, 959–968.

Sansinena, M., Santos, M. V., Zaritzky, N., Chirife, J. 2012. Comparison of heat transfer in liquid and slush nitrogen by numerical simulation of cooling rates for French straws used for sperm cryopreservation. Theriogenology, 77, 1717–1721.

Scheller, H. V. & Ulvskov, P. 2010. Hemicelluloses. Annual Review of Plant Biology, Vol 61, 61, 263–289.

Schneider, C. A., Rasband, W. S. & Eliceiri, K. W. 2012. NIH Image to ImageJ: 25 years of image analysis. Nature Methods, 9, 671–675.

Schweingruber, F. H. 2007. Wood structure and environment, Berlin, Springer.

Sczostak, A. 2009. Cotton Linters: An Alternative Cellulosic Raw Material. Macromolecular Symposia, 280, 45–53.

Simmons, T. J., Mortimer, J. C., Bernardinelli, O. D., Poppler, A. C., Brown, S. P., Deazevedo, E. R., Dupree, R. & Dupree, P. 2016. Folding of xylan onto cellulose fibrils in plant cell walls revealed by solid-state NMR. Nature Communications, 7.

Takenaka, Y., Watanabe, Y., Schuetz, M., Unda, F., Hill, J., Phookaew, P., Yoneda, A., Mansfield, S. D., Samuels, L., Ohtani, M & Demura, T. 2018. Patterned deposition of xylan and lignin is independent from that of the secondary wall cellulose of Arabidopsis xylem vessels. Plant Cell, 30, 2663–2676.

Taylor, N. G., Scheible, W. R., Cutler, S., Somerville, C. R. & Turner, S. R. 1999. The irregular xylem3 locus of arabidopsis encodes a cellulose synthase required for secondary cell wall synthesis. Plant Cell, 11, 769–779.

TEAM, R. C. 2014. R: A language and environment for statistical computing. [Online]. Vienna Austria. Available: http://www.R-project.org/ [Accessed].

Terashima, N., Kitano, K., Kojima, M., Yoshida, M., Yamamoto, H. & Westermark, U. 2009. Nanostructural assembly of cellulose, hemicellulose, and lignin in the middle layer of secondary wall of ginkgo tracheid. Journal of Wood Science, 55, 409–416.

Terrett, O. M., Dupree, P. 2019. Covalent interactions between lignin and hemicelluloses in plant secondary cell walls. Current Opinion in Biotechnology, 56, 97–104.

Thomas, L. H., Altaner, C. M. & Jarvis, M. C. 2013. Identifying multiple forms of lateral disorder in cellulose fibres. Journal of Applied Crystallography, 46, 972–979.

Thomas, L. H., Forsyth, V. T., Martel, A., Grillo, I., Altaner, C. M. & Jarvis, M. C. 2014. Structure and spacing of cellulose microfibrils in woody cell walls of dicots. Cellulose, 21, 3887–3895.

Thomas, L. H., Forsyth, V. T., Martel, A., Grillo, I., Altaner, C. M. & Jarvis, M. C. 2015. Diffraction evidence for the structure of cellulose microfibrils in bamboo, a model for grass and cereal celluloses. Bmc Plant Biology, 15.

Turner, S. & Kumar, M. 2017. Cellulose synthase complex organization and cellulose microfibril structure. Philosophical Transactions of the Royal Society A, 376.

Turner, S. R. & Somerville, C. R. 1997. Collapsed xylem phenotype of Arabidopsis identifies mutants deficient in cellulose deposition in the secondary cell wall. Plant Cell, 9, 689–701.

Vanholme, R., Demedts, B., Morreel, K., Ralph, J., Boerjan, W. 2010. Lignin biosynthesis and structure. Plant Physiology, 132, 1781–1789.

Vanholme, R., Storme, V., Vanholme, B., Sundin, L., Christensen, J. H., Goeminne, G., Halpin, C., Rohde, A., Morreel, K. & Boerjan, W. 2012. A Systems Biology View of Responses to Lignin Biosynthesis Perturbations in Arabidopsis. Plant Cell, 24, 3506–3529.

Whitney, S. E. C., Brigham, J. E., Darke, A. H., Reid, J. S. G. & Gidley, M. J. 1998. Structural aspects of the interaction of mannan-based polysaccharides with bacterial cellulose. Carbohydrate Research, 307, 299309.

Xiong, G., Cheng, K. & Pauly, M. Xylan O-Acetylation Impacts Xylem Development and Enzymatic Recalcitrance as Indicated by the Arabidopsis Mutant tbl29. Molecular Plant, 6, 1373–1375.

Yu, L., Lyczakowski, J. J., Pereira, C. S., Kotake, T., Yu, X., Li, A., Mogelsvang, S., Skaf, M. S. & Dupree, P. 2018. The patterned structure of galactoglucomannan suggests it may bind to cellulose in seed mucilage. Plant Physiology, 178, 1011–1026.

Zhang, T., Zheng, Y. Z. & Cosgrove, D. J. 2016. Spatial organization of cellulose microfibrils and matrix polysaccharides in primary plant cell walls as imaged by multichannel atomic force microscopy. Plant Journal, 85, 179–192.

Zhang, X. Y., Dominguez, P. G., Kumar, M., Bygdell, J., Miroshnichenko, S., Sundberg, B., Wingsle, G. & Niittyla, T. 2018. Cellulose Synthase Stoichiometry in Aspen Differs from Arabidopsis and Norway Spruce. Plant Physiology, 177, 1096–1107.

