## Supporting Information for "Structural imaging of native cryo-preserved secondary cell walls reveals presence of macrofibrils composed of cellulose, lignin and xylan"

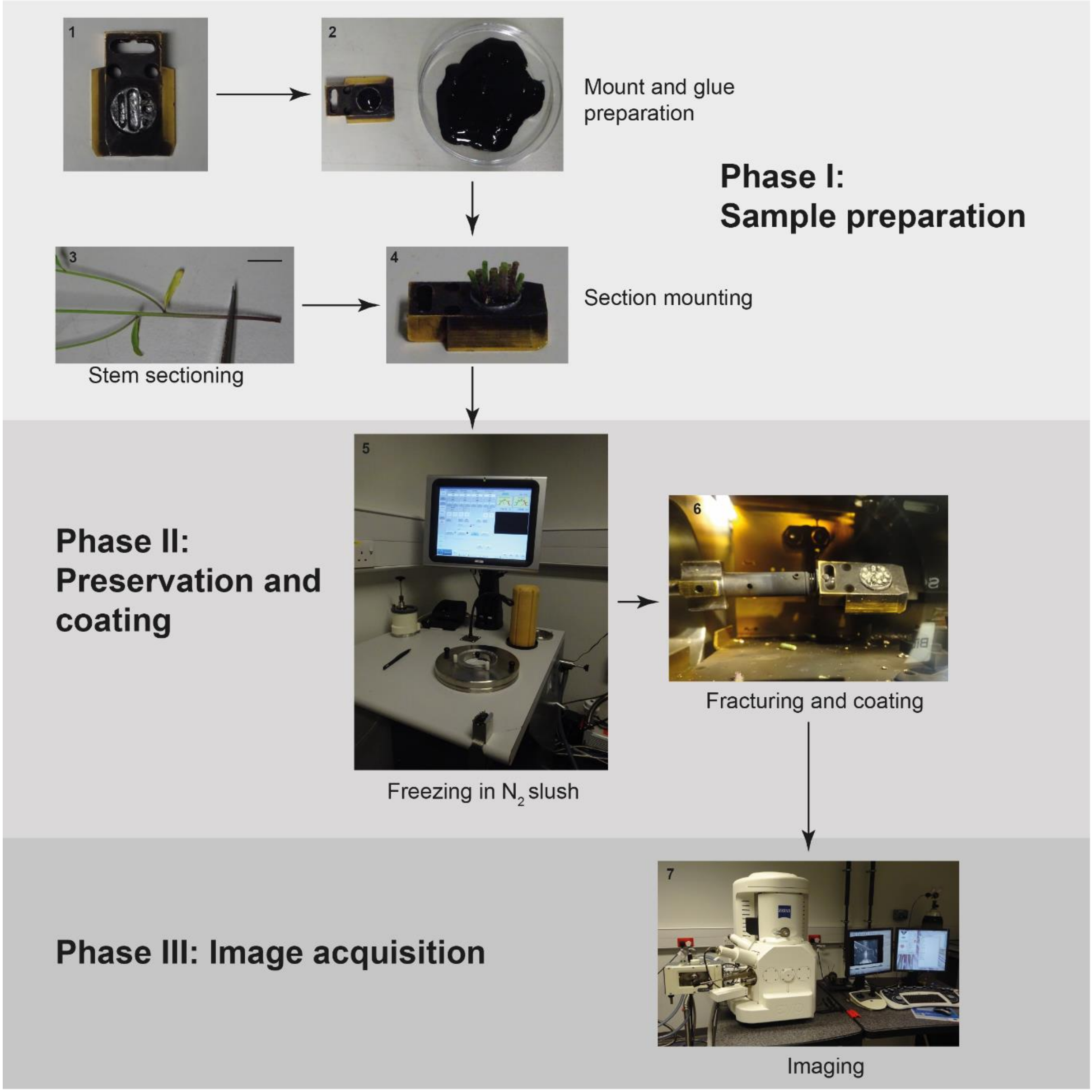


**Figure S1 Overview of the cryo-SEM procedure.** Presented images demonstrate sample preparation required for Arabidopsis imaging. Same protocol was applied for other samples analysed. Step 3 size bar is 1 cm long.

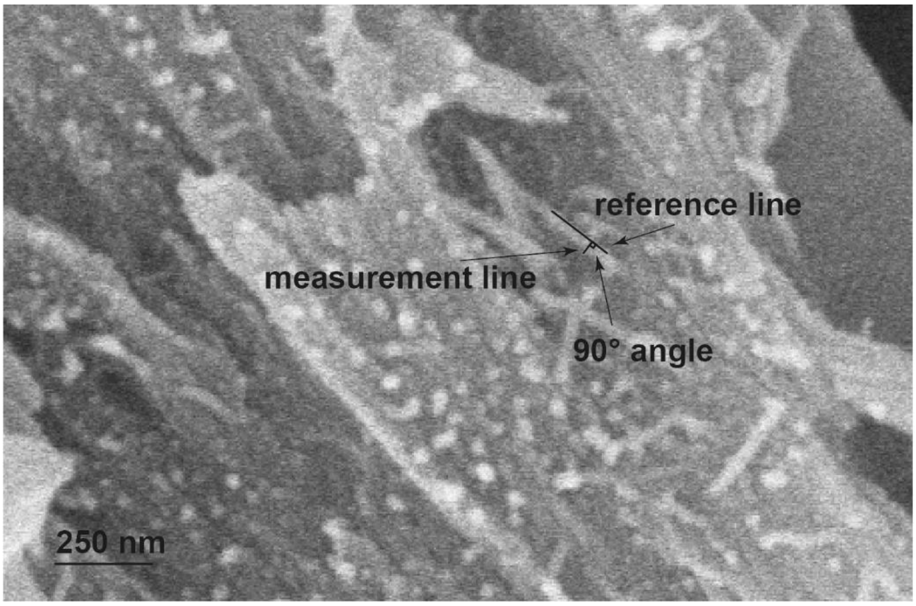


**Figure S2 Measurement of macrofibril diameter.** Example of Arabidopsis macrofibril measurement. A reference line and a perpendicular measurement line (equal in length to the fibril width) are marked.

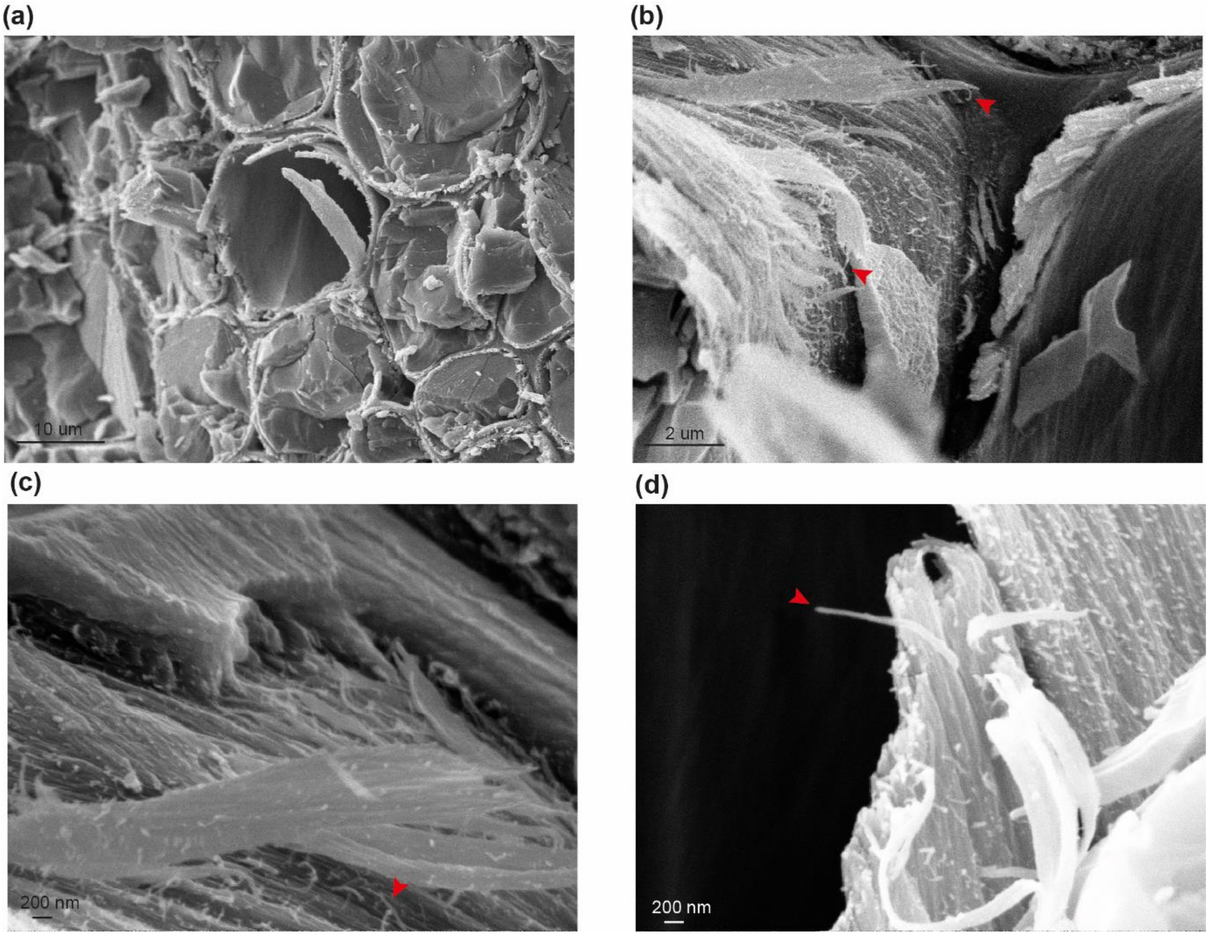


**Figure S3 Cryo-SEM analysis of Ginkgo cell walls.** (a) to (d) shows representative images at different magnification from analysis of stem sections from Ginkgo branches. Red arrows indicate macrofibrils. Size bar is provided for each image.

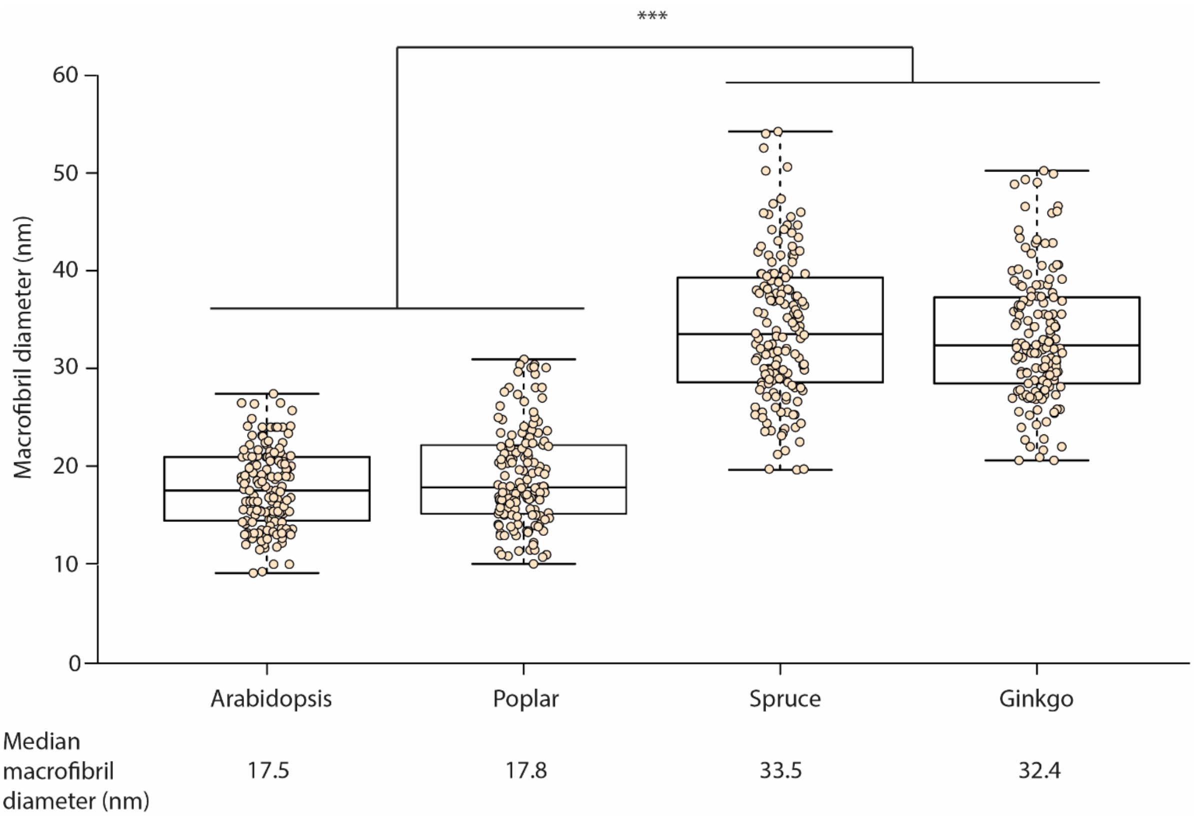


**Figure S4 Comparison of macrofibril diameter in Arabidopsis, poplar, spruce and Ginkgo.** N = 150 macrofibrils. Boxplots mark a median and show between 25th and 75th percentile of the data. *** denotes $p \leq 0.00001$ in Tukey test following ANOVA. No statistically significant difference was observed for the Arabidopsis- poplar and spruce-Ginkgo pairs.

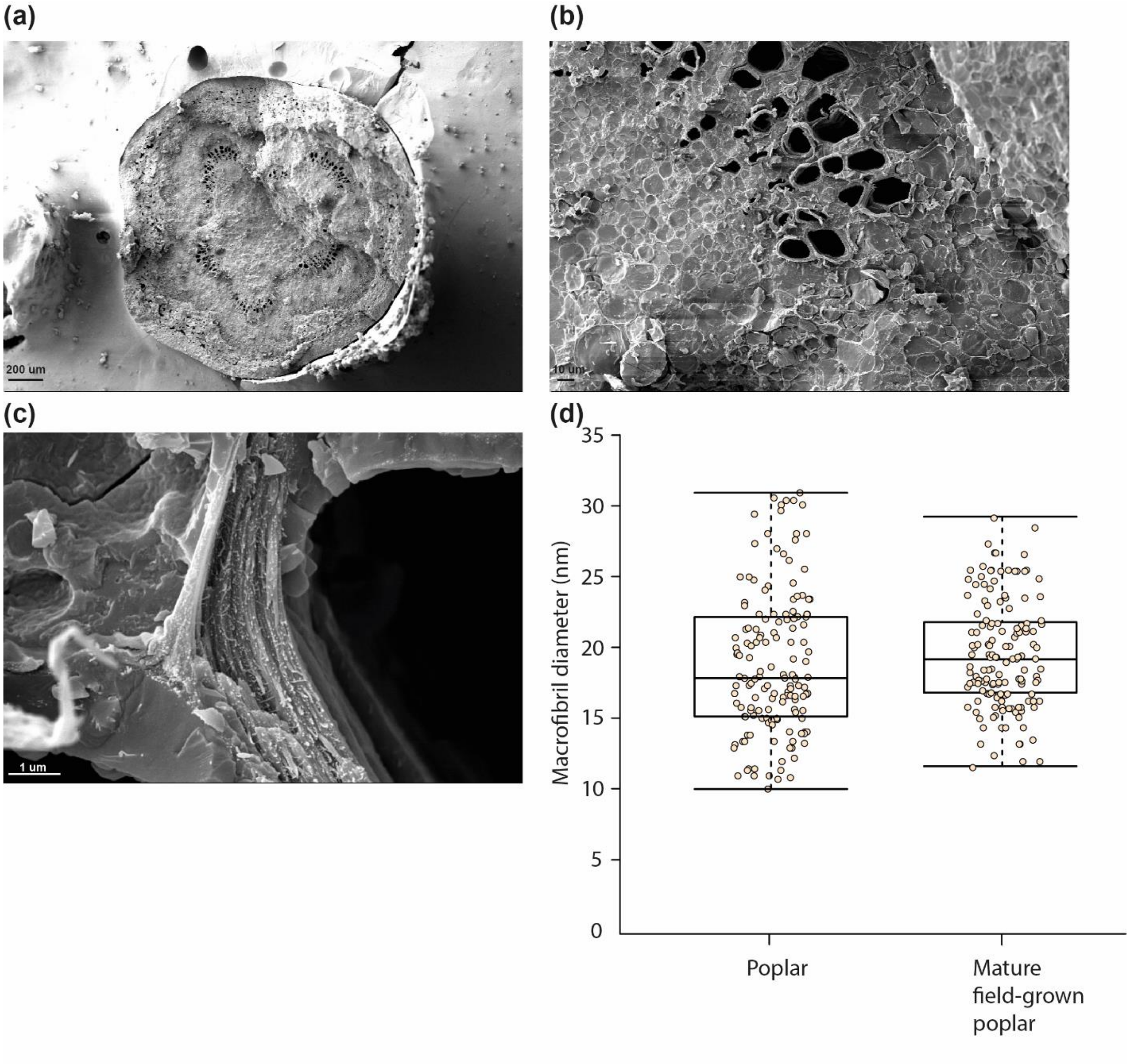


**Figure S5 Imaging of macrofibrils in field grown poplar.** (a)-(c) representative images of poplar (*Populus tremula*) hardwood from sample isolated from a mature tree grown at the Cambridge University Botanic Garden. Size bars are provided for each image. (d) Comparison of macrofibril diameter in *in vitro* and field grown poplar. N = 150 macrofibrils. Boxplots mark a median and show between 25th and 75th percentile of the data. No statistically significant difference was observed for the pair.

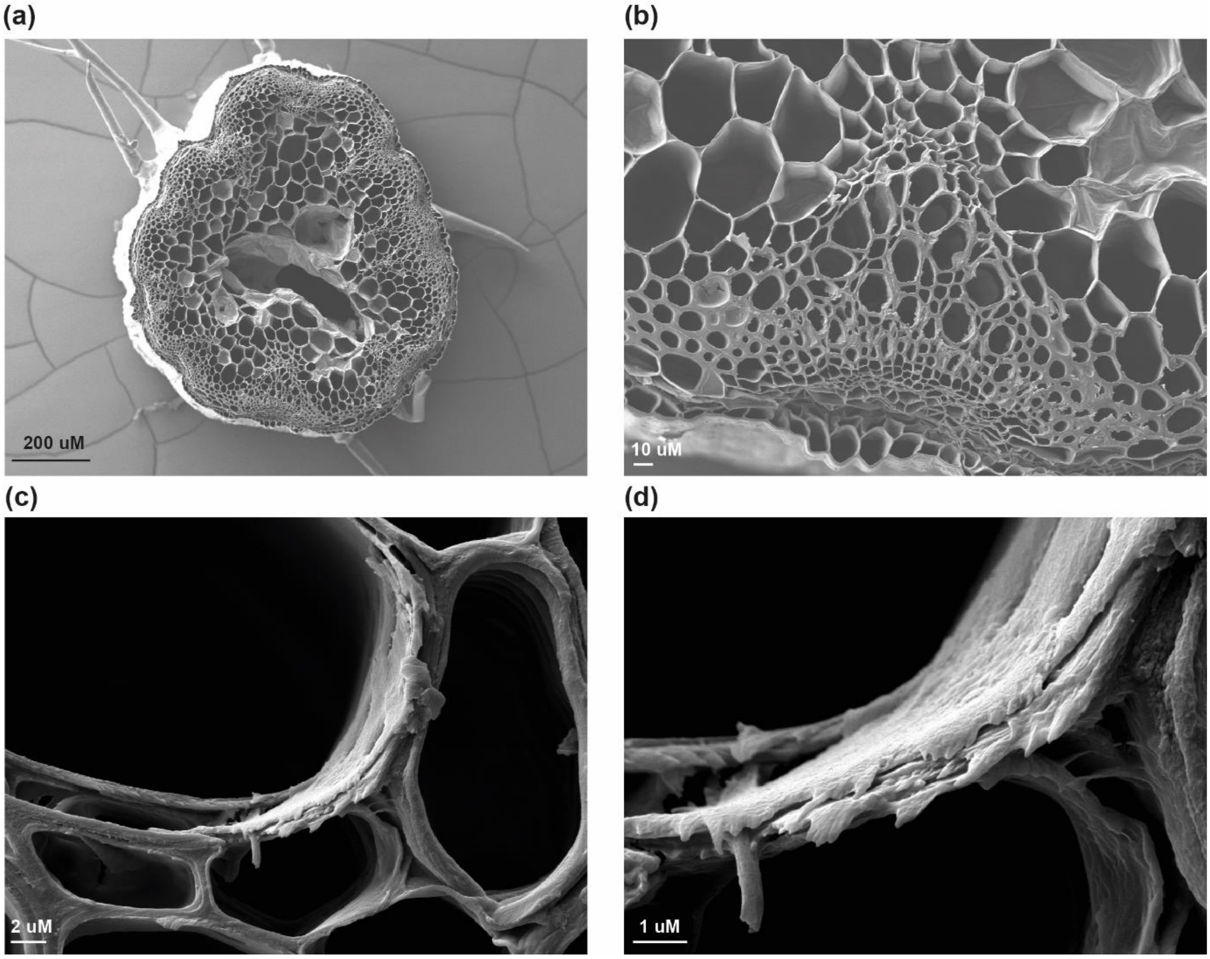


**Figure S6 Analysis of native Arabidopsis samples without the cryo-preservation protocol. (a)-(d)** Representative images of WT Arabidpsis stems imaged without the cryo-preservation protocol. Size bar is provided for each image.

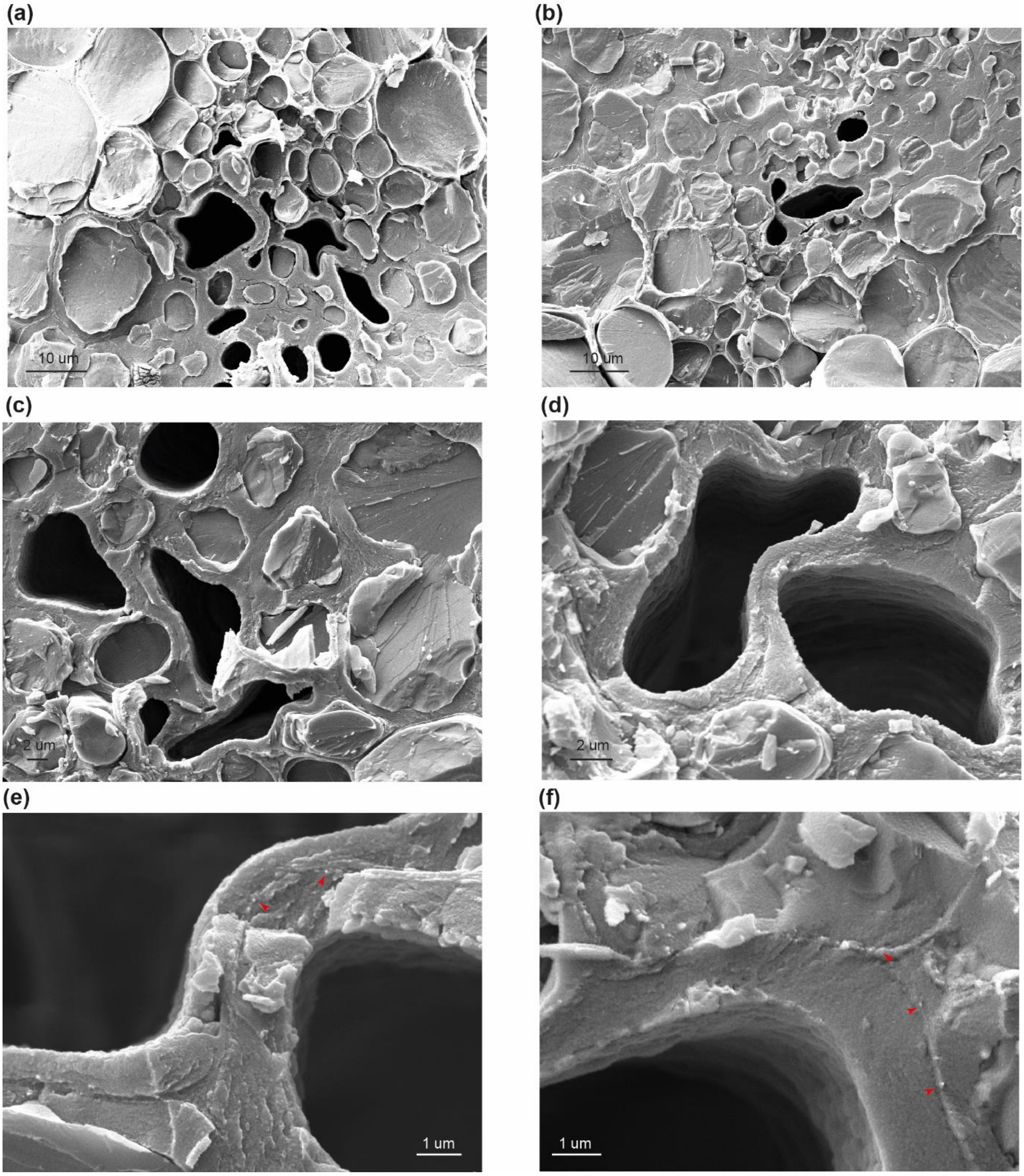


**Figure S7 Cryo-SEM analysis of vessel collapse and primary cell wall cellulose in *irx3* Arabidopsis plants.** (a) to (d) shows vessel collapse on representative images at different magnification from analysis of stem sections from *irx3* plants. Size bar is provided for each image. (e) and (f) show putative primary cell wall cellulose structures marked with red arrows.
